# CCR4-NOT subunit CCF-1/CNOT7 interacts with the PAL-1/CDX-1 transcription factor to regulate multiple stress responses in *Caenorhabditis elegans*

**DOI:** 10.1101/2022.06.13.495535

**Authors:** Hadi Tabarraei, Brandon M. Waddell, Kelly Raymond, Sydney M. Murray, Ying Wang, Keith P. Choe, Cheng-Wei Wu

**Author notes:** For correspondence (CWW).

## Abstract

CCR4-NOT is a versatile eukaryotic protein complex that controls multiple steps in gene expression regulation from synthesis to decay. In yeast, CCR4-NOT has been implicated in stress response regulation, though this function in other organisms remains unclear. In a genome-wide RNAi screen, we identified a subunit of the CCR4-NOT complex, *ccf-1*, as a requirement for the *C. elegans* transcriptional response to cadmium and acrylamide stress. Using whole-transcriptome RNA sequencing, we show that knockdown of *ccf-1* attenuates the activation of a broad range of stress protective genes in response to cadmium and acrylamide, including those encoding heat shock proteins and glutathione s-transferases. Consistently, survival assays show that knockdown of *ccf-1* decreases *C. elegans* stress resistance. A yeast-2-hybrid screen using a CCF-1 bait identified the homeobox transcription factor PAL-1 as a physical interactor. Knockdown of *pal-1* inhibits the activation of *ccf-1* dependent stress genes and reduces *C. elegans* stress resistance. Gene expression analysis reveals that knockdown of *pal-1* down-regulates the mRNA levels of *elt-2* and *elt-3*, which serves as the master transcriptional co-regulators of stress response in the *C. elegans* intestinal and epidermal tissues respectively. These results reveal a new role for CCR4-NOT in stress response regulation with PAL-1 through the transcriptional control of *elt-2* and *elt-3* in *C. elegans*.

## INTRODUCTION

In response to environmental stress, organisms can mount various adaptive responses to promote survival. At the cellular level, a major form of adaptive response is the transcriptional activation of genes that serve to prevent or repair cellular damage during periods of stress. Examples of such responses include induction of xenobiotic detoxification and antioxidant genes to alleviate oxidative stress or the activation of chaperone genes functioning in protein folding to maintain proteostasis under heat shock (Sarge et al. 1993; An et al. 2005). In the nematode *Caenorhabditis elegans*, various transcriptional pathways that regulate environmental stress response have also promoted longevity (Tullet et al. 2008; Morley & Morimoto 2004). Consistently, impaired stress response contributes to aging in *C. elegans* and has also been linked to various human diseases (Hekimi et al. 2011; Tullet et al. 2008; Morley & Morimoto 2004). As such, uncovering new insights toward understanding molecular mechanisms of stress response pathways are of widespread interest.

Our previous work in *C. elegans* showed that exposure to the heavy metal cadmium strongly activates a gene named *numr-1* (<u>nu</u>clear localized <u>m</u>etal <u>r</u>esponse), which encodes a putative RNA binding protein required for cadmium resistance (Wu et al. 2019). The mechanism of cadmium induced *numr-1* activation appears to be distinct from the hallmark induction of metallothionein encoding genes that maintain homeostasis by scavenging and removing xenobiotic metals (Sabolić et al. 2010). Cadmium activation of *numr-1* partially signals through the transcription factor HSF-1 (<u>H</u>eat <u>S</u>hock <u>F</u>actor), whereas metallothionein gene activation in *C. elegans* requires the DAF-16 (<u>DA</u>uer <u>F</u>ormation)/FoxO transcription factor (Barsyte et al. 2002; Wu et al. 2019). Given that cadmium is known to induce many types of cellular alterations including oxidative stress, DNA damage, and RNA splicing disruption, it is not surprising that multiple stress response pathways are simultaneously activated to promote cellular resistance (Bertin & Averbeck 2006; Chomyshen et al. 2022). Additionally, cadmium activates the expression of genes encoding various detoxification enzymes involved in glutathione metabolism that is controlled by the SKN-1 (<u>SK</u>i<u>N</u>head)/Nrf2 transcription factor (An et al. 2005). A recent study has also implicated a role for the nuclear hormone receptor HIZR-1 (<u>HI</u>gh <u>Z</u>inc activated nuclear <u>R</u>eceptor) in regulating the expression of cadmium and zinc responsive genes in cooperation with the mediator transcriptional complex (Shomer et al. 2019). The involvement of multiple transcription factors in response to metal stress in *C. elegans* may also reflect the lack of the MTF-1 (<u>M</u>etal responsive <u>T</u>ranscription <u>F</u>actor) gene in nematodes that serves as the conserved regulator of metal induced transcriptional response in fly, fish, and mammals (Günther et al. 2012). While the requirement of multiple transcription factors involved in stress responses are well recognized, less known are co-regulators that assist in these transcriptional activation events.

In this study, we used *numr-1* as a stress marker to identify a new role for the CCR4-NOT (<u>C</u>arbon <u>C</u>atabolite <u>R</u>epression <u>4</u> – <u>N</u>egative <u>O</u>n <u>T</u>ATA-less) complex in regulating *C. elegans* transcriptional response to multiple environmental stress. The CCR4-NOT complex was originally identified as an eukaryotic deadenylase that post-transcriptionally regulates mRNA stability through poly-A tail removal (Nousch et al. 2013; Collart 2016). Subsequent studies in yeast have also revealed a role for the CCR4-NOT in transcriptional initiation through interactions with RNA polymerase II, implicating a broad regulatory function of this complex in mRNA synthesis and decay (Collart 2016). A direct role for CCR4-NOT complex in stress regulation has also been recently reported in yeast (Mulder et al. 2005). Here, we demonstrate that the *C. elegans* CCR4-NOT deadenylase subunit *ccf-1* (<u>CC</u>R4 associated <u>f</u>actor) is broadly required for the stress induced transcriptional programming in *C. elegans* in response to cadmium and acrylamide. Our results show that *ccf-1* is essential for normal aging along with stress resistance, and its knockdown also shortens the lifespan of multiple long-lived mutants. Through the yeast-2 hybrid (Y2H) system, we identified the homeobox PAL-1 (<u>P</u>osterior <u>AL</u>ae) transcription factor as a novel physical interactor of CCF-1 and uncovered a previously undescribed role for this protein in the regulation of stress response genes in *C. elegans.* Overall, this study provides new insights into genetic regulators of transcriptional response and resistance to environmental stress in *C. elegans*.

## RESULTS

### Components of the CCR4-NOT complex are necessary for the cadmium stress response

We previously showed that the cadmium inducible *numr-1* gene encodes as a putative RNA binding protein that influences RNA splicing in *C. elegans* (Wu et al. 2019). To screen for regulators of this stress response, we performed a genome-wide RNAi screen using a *numr-1p*::GFP transcriptional reporter to identify genes that when knocked down inhibit the induction of *numr-1p*::GFP after cadmium exposure (**Figures 1a-b**). We identified 49 genes that when knocked down reduced *numr-1p*::GFP after cadmium and had minimal effect on *C. elegans* development. Enrichment analysis of these 49 genes revealed functional annotation to two protein complexes, the CCR4-NOT complex and the Arp2/3 (<u>A</u>ctin-related <u>p</u>rotein) complex (**Figure 1c; Table S1**). Fluorescence levels of *numr-1p*::GFP after cadmium exposure were highly suppressed when four genes encoding subunits of the CCR4-NOT complex and three genes encoding subunits of the Arp2/3 complex were knocked down by RNAi (**Figure 1c-d**). The CCR4-NOT complex is well characterized for its role as an mRNA deadenylase that facilitates poly-A tail removal, a cellular process that decreases transcript stability that is highly conserved in yeast, worm, and human (Shirai et al. 2014; Nousch et al. 2013). However, emerging evidence in yeast suggests that the CCR4-NOT complex is also involved in transcriptional regulation, though this molecular mechanism is less understood (Mulder et al. 2005; Kruk et al. 2011). As such, we focused on characterizing the role of the CCR4-NOT complex in the regulation of stress induced transcription in this study.

**Figure 1.**
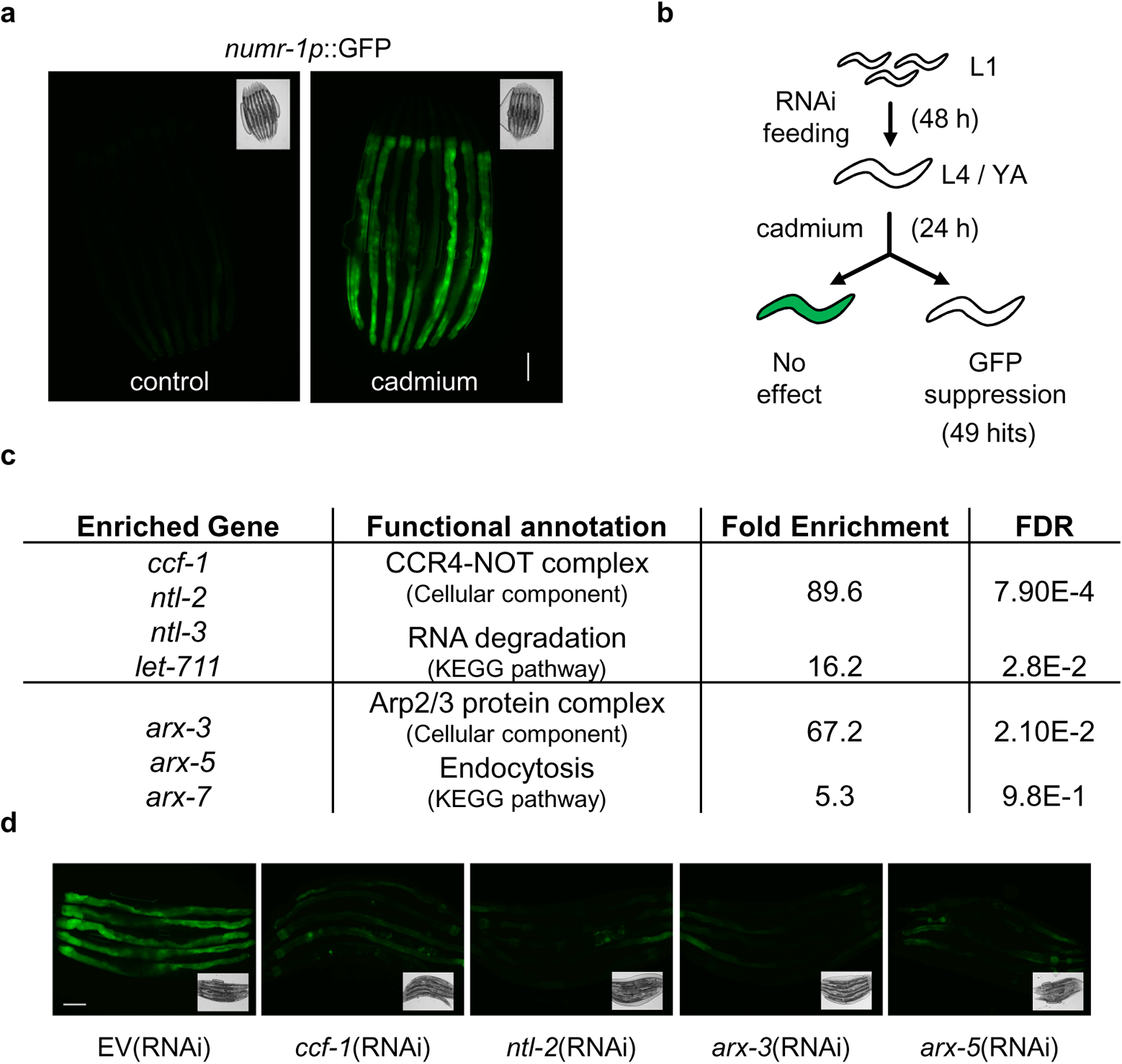
Genome-wide RNAi screen reveals a requirement for genes encoding the CCR4-NOT complex for cadmium induced activation of *numr-1p*::GFP. **a.** Representative fluorescence and brightfield micrographs of worms expressing *numr-1p*::GFP under control and cadmium conditions. **b**. Workflow of genome-wide RNAi screen. **c**. DAVID enrichment of dsRNA clones that inhibited cadmium induced *numr-1p*::GFP expression. **d**. Representative fluorescent and brightfield micrographs of worms fed dsRNA resulting in the inhibition of cadmium induced expression of *numr-1p*::GFP. The scale bar is 100 μm.

To determine if the CCR4-NOT complex is required for the activation of a broad range of cadmium inducible genes, we used RNAi to knock down the *ccf-1/CNOT7* gene that encodes the deadenylase subunit of the CCR4-NOT complex (Nousch et al. 2013). Using qPCR, we found that expressions of various classes of cadmium induced genes are significantly reduced in worms fed with *ccf-1* RNAi as compared to the empty vector (EV) control after cadmium exposure, including those encoding glutathione s-transferase (*gst*) and heat shock protein (*hsp*) genes (**Figure 2a**). Consistent with *ccf-1* functioning as a requirement for the cadmium stress response, knockdown of *ccf-1* reduced *C. elegans* cadmium resistance; meanwhile, knockdown of *ccf-1* also significantly reduced *C. elegans* lifespan, suggesting that *ccf-1* is required for normal aging (**Figure 2b**). To determine if other subunits of the CCR4-NOT complex are also required for the activation of cadmium responsive genes (**Figure 2c**), we knocked down three other genes identified from the genome-wide RNAi screen that encode subunits of the CCR4-NOT complex and performed qPCR analysis. RNAi knockdown of *ntl-2* (<u>N</u>O<u>T</u>-<u>l</u>ike)/*CNOT2* suppressed the activation of nine out of ten cadmium induced genes tested, whereas knockdown of *ntl-3*/*CNOT3* and *let-711* (<u>LET</u>hal)/*CNOT1* suppressed the activation of two and five out of ten cadmium induced genes respectively (**Figure 2d**). The relative variance in gene expression levels may suggest a differential requirement of each CCR4-NOT subunit in regulating cadmium induced transcriptional response. It is also possible that the difference may be due to varying degrees of RNAi penetrance between the dsRNA clones. Although, RNAi depletion of *ccf-1*, *ntl-2,* and *ntl-3* all led to 100% F_1_ embryonic lethality and *let-711* RNAi led to P_0_ L3/L4 arrest (data not shown). Overall, these data suggest that subunits of the CCR4-NOT complex are required for *C. elegans* transcriptional response to cadmium stress and that the *ccf-1* deadenylase subunit is essential for normal aging and cadmium survival.

**Figure 2.**
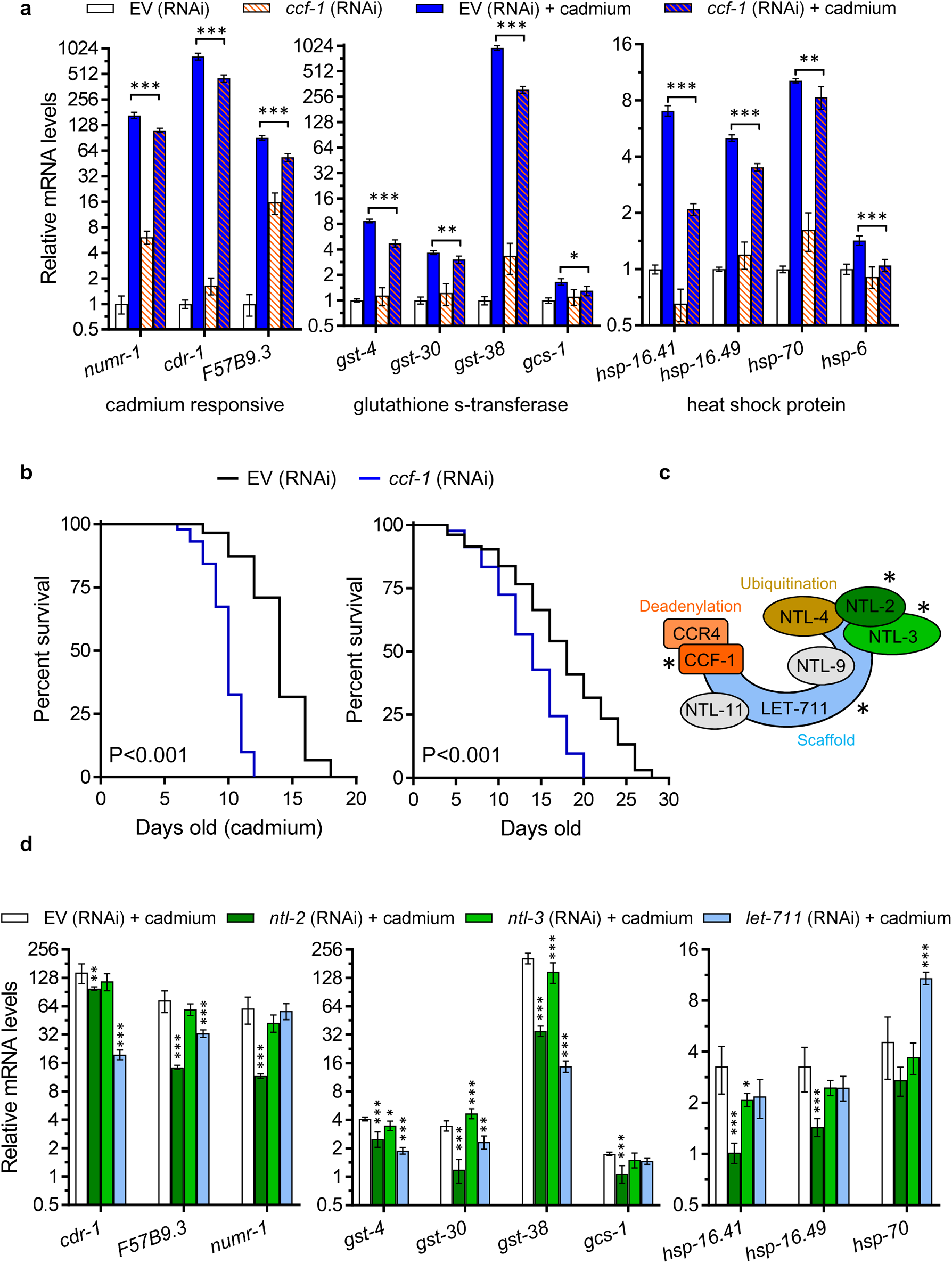
Components of the CCR4-NOT complex regulate the cadmium stress response. **a.** Relative expression of cadmium responsive genes in N2 worms fed with EV or *ccf-1* RNAi under basal conditions or after exposure to cadmium. **b**. Cadmium survival and lifespan curves of N2 worms fed with EV or *ccf-1* RNAi, P<0.001 as determined by the log-rank test. Results for trial 1 are shown for cadmium survival and trial 2 for lifespan curve, full results and statistics for all trials are presented in Table S2. **c.** Components of the *C. elegans* CCR4-NOT complex with * indicating genes identified to be required for cadmium induced *numr-1p*::GFP. **d.** Fold change in expression of cadmium responsive genes in N2 worms fed with EV, *ntl-2*, *ntl-3*, or *let-711* RNAi after cadmium exposure relative to basal condition. For **a** and **d**, bar graphs display means and error bars indicate the standard deviation of N = 4 samples. *P<0.05, **P<0.01, and ***P<0.001 compared to EV (RNAi) + cadmium as determined by two-way ANOVA with Holm-Sidak *post hoc* tests.

### *ccf-1* is required for the transcriptional response activated by multiple stressors

Given that knockdown of *ccf-1* led to a consistent reduction in expression of select cadmium inducible genes, we next examined the effect of *ccf-1* knockdown on the whole transcriptome before and after cadmium exposure using RNA sequencing (**Figure 3a**). Knockdown of *ccf-1* under basal conditions led to differential expression of 3,901 genes by more than 2-fold, with the majority of these genes up-regulated (**Figure 3b**). This is consistent with the known deadenylase function of CCF-1 where its knockdown results in the retention of mRNA poly-A tail that increases transcript stability (Nousch et al. 2013). Enrichment analysis revealed that genes up-regulated after *ccf-1* knockdown cluster to KEGG pathways including ABC transporters, ribosome biogenesis, and lipid metabolism, whereas genes down-regulated by *ccf-1* RNAi cluster to the lysosome pathway (**Figure 3b**).

**Figure 3.**
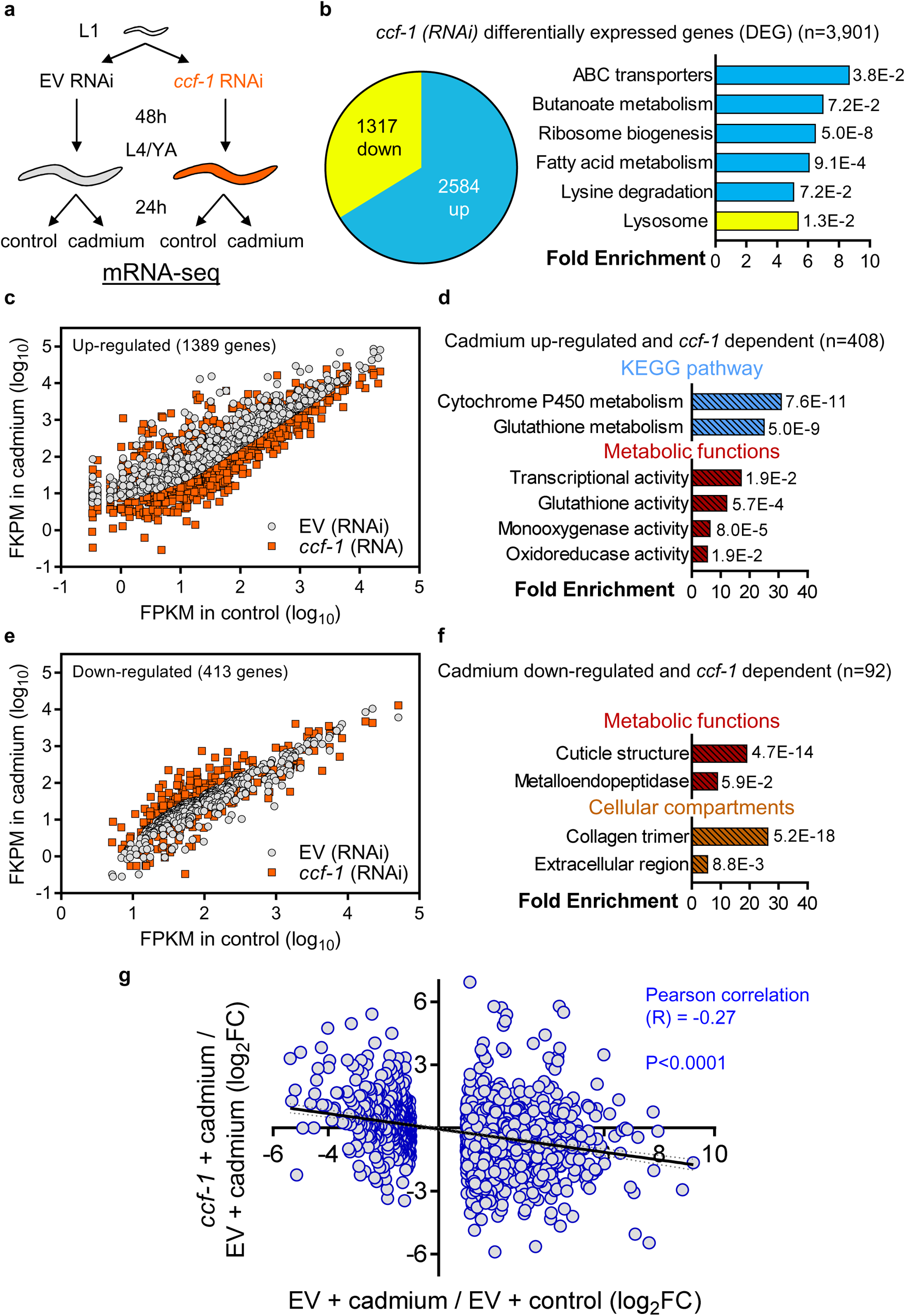
Whole-transcriptome sequencing reveals *ccf-1* is broadly required for cadmium induced gene expression. **a**. Experimental workflow used for preparing RNA-sequencing samples. **b**. Pie chart illustrating the number of genes significantly up-regulated or down-regulated in N2 worms by > 2-fold after *ccf-1* (RNAi) compared to EV. The bar graph illustrates DAVID analysis of KEGG pathways significantly enriched for genes up-regulated (blue) or down-regulated (yellow) after *ccf-1* (RNAi) compared to EV. Fold enrichment and FDR values are shown for each KEGG pathway. **c.** Scatter-plot of 1,389 cadmium up-regulated genes (>2-fold) in N2 worms fed with EV compared to their corresponding expression in worms fed with *ccf-1* (RNAi). **d**. Enriched KEGG pathways and metabolic functions of 408 *ccf-1* dependent cadmium up-regulated genes. **e.** Scatter-plot of 413 cadmium down-regulated genes (>2-fold) in N2 worms fed with EV compared to their corresponding expression in N2 worms fed with *ccf-1* (RNAi). **f**. Enriched metabolic functions and cellular compartments of 92 *ccf-1* dependent cadmium down-regulated genes. **g.** Linear regression analysis of log_2_ fold change between *ccf-1* (RNAi) and EV (plotted on the y-axis) for the 1,802 genes up or down-regulated by cadmium in N2 worms fed with EV (plotted on the x-axis). P<0.001 as determined by the F-test.

Next, we examined the effects of *ccf-1* knockdown on genes that are differentially regulated by cadmium more than 2-fold. Scatterplot analysis of the 1,389 genes up-regulated by cadmium shows that their expressions are generally reduced after *ccf-1* knockdown in comparison to EV RNAi. Of the 1,389 cadmium up-regulated genes, 408 were found to be reduced by >2-fold in *ccf-1* knocked down worms indicating their dependence on *ccf-1* for cadmium induced gene expression (**Figure 3c**). Enrichment analysis of the 408 *ccf-1* dependent cadmium genes reveal that they cluster to xenobiotic detoxification pathways and metabolic functions including cytochrome P450 metabolism and glutathione activity (**Figure 3d**). We then examined how *ccf-1* knockdown influences cadmium down-regulated genes and found that their expressions are generally increased after *ccf-1* knockdown in comparison to EV RNAi (**Figure 3e**). Enrichment analysis of the 92 *ccf-1* dependent cadmium down-regulated genes reveals they function within cuticle structure and collagen/extracellular maintenance (**Figure 3f**; no enrichment towards KEGG pathway was found). The broad requirement for *ccf-1* in cadmium induced gene expression is further illustrated through linear regression analysis which revealed a Pearson correlation (R) value of −0.27 (P<0.001) for relative fold change between *ccf-1* RNAi and EV RNAi for the 1,802 genes differentially expressed by cadmium (**Figure 3g**). These results demonstrate that *ccf-1* is an essential factor in mediating cadmium induced transcriptional programming.

As *ccf-1* is required for the expression of antioxidant genes in response to cadmium, we then examined if *ccf-1* is required for mounting a stress response to acrylamide, which is a potent chemical inducer of oxidative stress (Wu et al. 2017). Using qPCR, we found that knockdown of *ccf-1* reduced the activation glutathione related antioxidant genes after acrylamide exposure including *gst-12*, *gst-30*, *gst-38*, and *gcs-1* (**Figure 4a**). Next, we expanded this analysis to the whole transcriptome by using RNA sequencing and found that genes differentially regulated by acrylamide are also broadly dependent on *ccf-1* (**Figure 4b-c**). Linear regression analysis of the relative fold change between *ccf-1* RNAi and EV RNAi for the 851 genes differentially expressed by acrylamide (>2-fold) showed a negative R-value of −0.32 (P<0.001) (**Figure 4d**). These results support a general requirement for *ccf-1* in the transcriptome regulation of the acrylamide stress response. Enrichment analysis of 220 *ccf-1* dependent acrylamide up-regulated genes cluster to KEGG pathways involved in cytochrome P450 and glutathione metabolism (**Figure 4e**). Consistent with *ccf-1* as a key regulator for the acrylamide stress response, knockdown of *ccf-1* reduced *C. elegans* survival to acrylamide (**Figure 4f**) and exacerbated acrylamide’s neurotoxicity toward dopaminergic neurons (**Figure 4g**).

**Figure 4.**
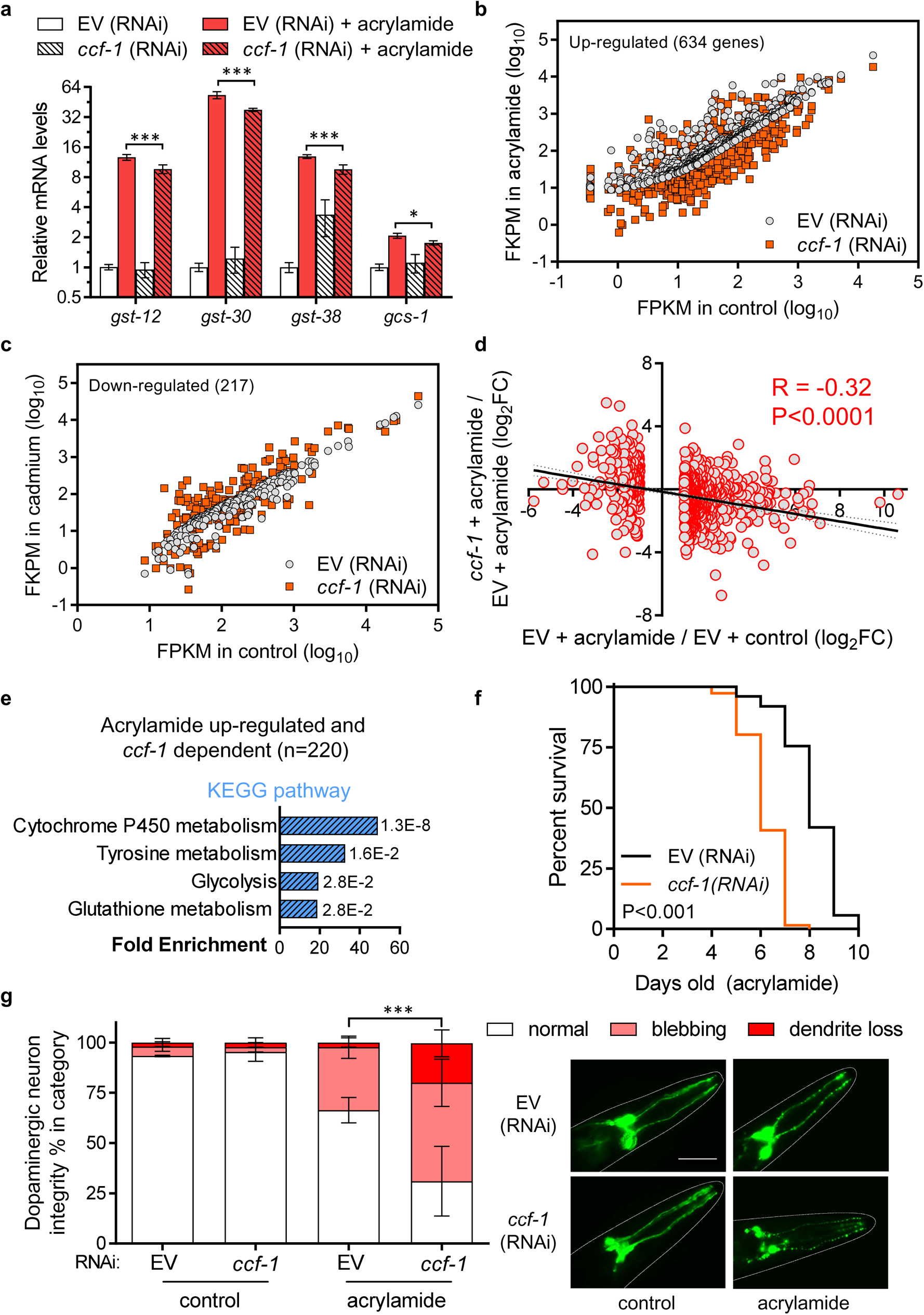
Depletion of *ccf-1* diminishes *C. elegans* resistance to acrylamide. **a.** Relative expression of *gst* genes and *gcs-1* in worms fed with EV or *ccf-1* (RNAi) under basal conditions, or after exposure to acrylamide. Bar graphs display means and error bars indicate standard deviation of N = 4 samples. *P<0.05 and ***P<0.001 compared to EV (RNAi) + acrylamide as determined by two-way ANOVA with Holm-Sidak *post hoc* tests. **b.** Scatter-plot of 637 acrylamide up-regulated and **c.** 217 acrylamide down-regulated genes in worms fed with EV compared to their corresponding gene expression in worms fed with *ccf-1* (RNAi). **d.** Linear regression analysis of log_2_ fold change between *ccf-1* (RNAi) and EV (plotted on the y-axis) for the 851 genes up or down-regulated by acrylamide in worms fed with EV (plotted on the x-axis). P<0.001 as determined by the F-test **e.** Enriched KEGG pathways of the 220 *ccf-1* dependent acrylamide up-regulated genes. **f.** Acrylamide survival curve of N2 worms fed with EV or *ccf-1* (RNAi), P<0.001 as determined by the log-rank test. Results for trial 2 are shown for acrylamide survival, full results and statistics for all trials are presented in Table S2. **g.** Dopaminergic neuron integrity of worms fed with EV or *ccf-1* (RNAi) under basal conditions, or treated with acrylamide. Results are combined from three trials of N = 15 worms per condition scored each trial. Representative fluorescent micrographs of control or acrylamide treated worms expressing the *dat-1p*::GFP reporter fed with EV or *ccf-1* (RNAi), scale bar = 50 µm. Error bars indicate standard error of the mean, ***P<0.001 as determined by the Chi-square test.

Additionally, we show that knockdown of *ccf-1* attenuates the activation of *gpdh-1p*::GFP in response to osmotic stress (**Figure S1a**) and reduced the expression of *hsp* genes in response to heat stress (**Figure S1b**). Overall, these results demonstrate a central role for *ccf-1* in mounting a transcriptome response to various environmental stressors and show that *ccf-1* is essential for *C. elegans* survival to cadmium and acrylamide stress.

### Stress resistant long-lived mutants required *ccf-1* for longevity

Transcriptomic analysis revealed that *ccf-1* is required for the expression of various stress responsive genes controlled by the transcription factors SKN-1 and HSF-1 that also promote longevity in *C. elegans* (Tullet et al. 2008; Baird et al. 2014). To test whether *ccf-1* is required for SKN-1 and HSF-1 dependent longevity, we knocked down *ccf-1* in the *skn-1(k1023)* gain of function mutant and a full-length *hsf-1(FL)* overexpression strain (Baird et al. 2014; Tang & Choe 2015). Knockdown of *ccf-1* completely suppressed the longevity phenotype of the *skn-1(k1023)* mutant (**Figure S2a**). Meanwhile, *hsf-1(FL)* worms fed with *ccf-1* RNAi lived on averaged +10% longer than N2 worms fed with *ccf-1* RNAi, this is reduced from an average of +28% in lifespan observed in EV RNAi fed worms suggesting that the longevity of *hsf-1(FL)* mutant is partially dependent on *ccf-1* (**Figure S2b**). The *skn-1* gene also acts downstream of the insulin signaling pathway to promote the longevity of the insulin receptor *daf-2* mutant (Tullet et al. 2008). Knockdown of *ccf-1* in the long-lived *daf-2(e1037)* mutants also significantly reduced its lifespan compared to EV RNAi, although the relatively lifespan extension of the *daf-2* mutant compared to N2 was similar when both strains were fed with *ccf-1* RNAi (average +216%) or EV RNAi (averaged +230%) (**Figure S2c**). Overall, these results show that *ccf-1* is required for the longevity of the *skn-1(k1023)* mutant and partially required for the longevity of the *hsf-1(FL)* and *daf-2(e1037)* mutants.

### CCF-1 localizes to the nucleus and interacts with the PAL-1 transcription factor

To gain insights into how *ccf-1* influences stress responsive genes, we generated a strain of *C. elegans* expressing an integrated *ccf-1p*::CCF-1::GFP transgene to test if CCF-1 protein expression is influenced by stress. CCF-1::GFP is expressed in multiple tissues and intestinal nuclear CCF-1::GFP can be observed in ~50% of the animals under basal conditions (**Figure S3a**). Exposure to cadmium or acrylamide did not alter the relative nuclear distribution or total fluorescence of CCF-1::GFP, suggesting that the expression and localization of the CCF-1 protein are not influenced by stress (**Figure S3b-c**). However, the nuclear presence of CCF-1 suggests potential alternative functions other than its characterized cytoplasmic deadenylase role.

Next, we employed the Y2H system using a GAL4-DBD-CCF-1 fusion protein to probe against a mixed-stage *C. elegans* GAL4-AD cDNA prey library to identify potential CCF-1 binding partners. The Y2H screen revealed two consistent prey binding partners of CCF-1 that corresponded to clones encoding the CCR4 and PAL-1 proteins (**Figure 5a, Figure S4**). CCR4 encodes a catalytic subunit of the CCR4-NOT complex and this protein has previously been shown to directly interact with CCF-1 in *C. elegans* (Nousch et al. 2013). Meanwhile, PAL-1 encodes the orthologue of the human caudal-related homeobox 1 (CDX1) transcription factor that is characterized by its role in embryonic posterior patterning and male tail development in *C. elegans* (Edgar et al. 2001). To test if PAL-1 co-regulates *ccf-1* dependent stress genes, we used RNAi to knock down *pal-1* and performed qPCR to quantify its effects on cadmium and acrylamide induced gene expressions. Knockdown of *pal-1* in the N2 wildtype strain did not lead to substantial changes to cadmium and acrylamide induced gene expression (data not shown), perhaps due to the low penetrance of the *pal-1* RNAi. Given that a null mutant is not available, we then repeated the experiment using the RNAi sensitive *rrf-3(pk1426)* strain where *pal-1* knockdown broadly reduced the activation of *ccf-1* dependent cadmium induced genes. (**Figure 5b**). Knockdown of *pal-1* in the *rrf-3(pk1426)* strain also reduced cadmium survival compared to the EV RNAi, suggesting a role for *pal-1* in stress resistance (**Figure 5c**). Similarly, knockdown of *pal-1* in the *rrf-3(pk1426)* worms treated with acrylamide attenuated the activation of *ccf-1* dependent *gst* genes and reduced acrylamide survival when compared to EV RNAi (**Figure 5d-e**).

**Figure 5.**
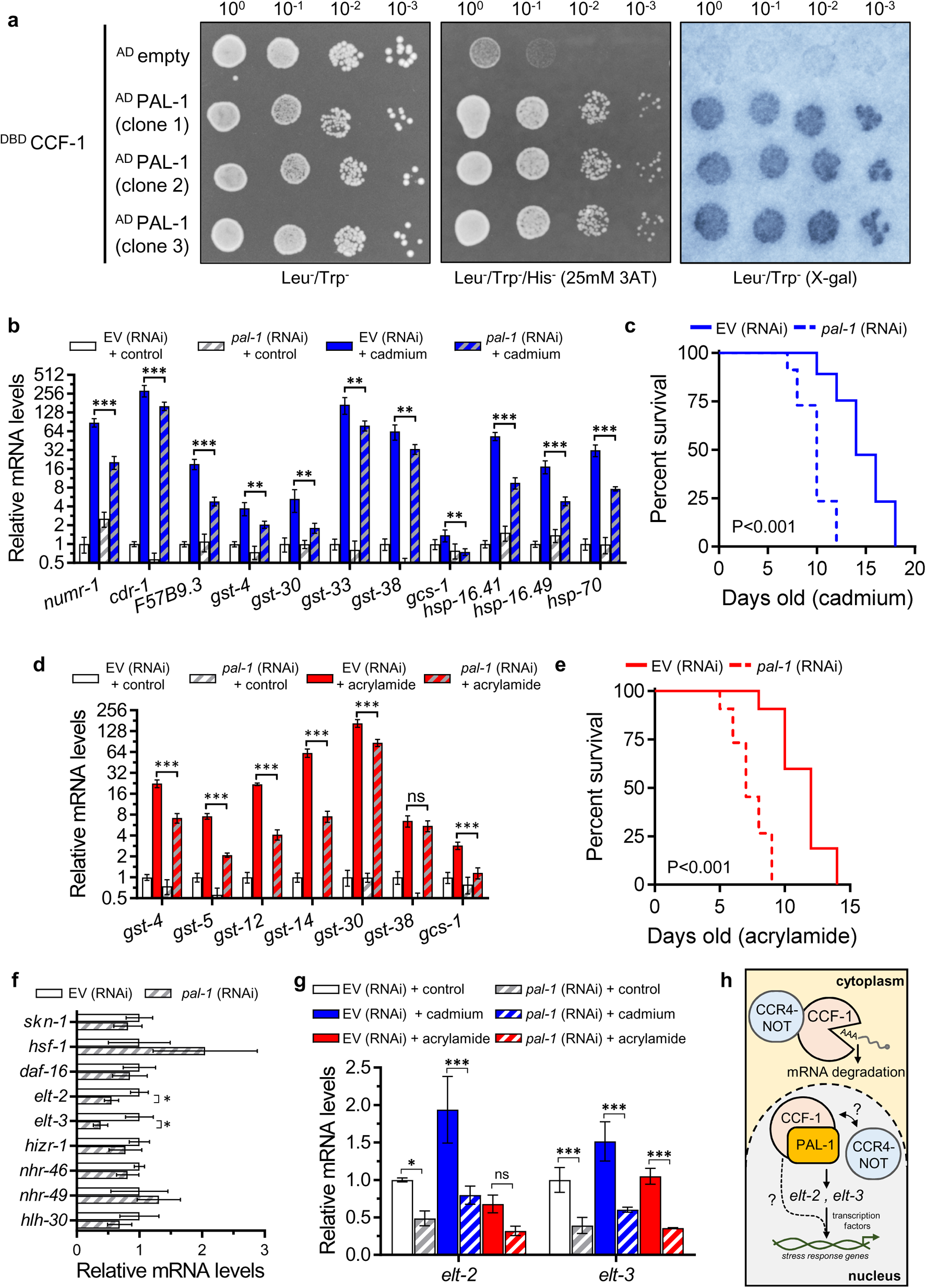
PAL-1 interacts with CCF-1 and regulates stress responsive genes. **a.** Y2H interactions between CCF-1 (bait) and three independent clones of PAL-1 (prey) on the control Leu^−^/Trp^−^ plate, Leu^−^/Trp^−^/His^−^/25 mM 3AT selection plate, and Leu^−^/Trp^−^ X-gal assay. Color in the X-gal image is artificially shaded blue. **b.** Relative expression of cadmium response genes under control or cadmium exposure in *rrf-3(pk1426)* mutant worms fed with EV or *pal-1* (RNAi). **c.** Cadmium survival of *rrf-3(pk1426)* worms fed with EV or *pal-1* (RNAi). **d.** Relative expression of acrylamide response genes under control or acrylamide exposure in *rrf-3(pk1426)* worms fed with EV or *pal-1* (RNAi). **e**. Acrylamide survival of *rrf-3(pk1426)* worms fed with EV or *pal-1* (RNAi). **f.** Relative expression of stress regulating transcription factors in *rrf-3(pk1426)* worms fed with EV or *pal-1* (RNAi). **g.** Relative expression of *elt-2* and *elt-3* mRNA levels under control, cadmium, or acrylamide exposure in *rrf-3(pk1426)* worms fed with EV or *pal-1* (RNAi). All bar graphs display means and error bars indicate a standard deviation of N = 4 samples. *P<0.05, **P<0.01, and ***P<0.001 as determined by two-way ANOVA with Holm-Sidak *post hoc* tests in **b**, **d**, and **g**. The student’s t-test corrected with Holm-Sidak multiple testing was used in **f** with *P<0.05. Trial 2 is shown for the survival assays in **c** and **e**, full results and statistics for all trials are presented in Table S2. **h**. Proposed mechanism of CCF-1 and PAL-1 function in *C. elegans* stress response regulation.

Previous studies in *C. elegans* have implicated a role for *pal-1* in embryogenesis through regulating the expression of key transcription factors such as HLH-1 (Helix Loop Helix) that controls body wall muscle development (Lei et al. 2009; Baugh et al. 2005). To test the hypothesis that *pal-1* may regulate the mRNA expression of stress responsive transcription factors, we knocked down *pal-1* in the *rrf-3(pk1426)* strain to determine its effect on nine transcription factors involved in stress response including *skn-1, hsf-1, daf-16, elt-2* (<u>e</u>rythroid-like transcription factor)*, elt-3*, *hizr-1, nhr-46* (<u>n</u>uclear <u>h</u>ormone receptor)*, nhr-49,* and *hlh-30* (An & Blackwell 2003; Guisbert et al. 2013; Lin et al. 2018; Budovskaya et al. 2008; Shomer et al. 2019; Goh et al. 2014). Of the nine transcription factors measured, we found that knockdown of *pal-1* led to a substantial decrease in both *elt-2* and *elt-3* mRNA levels by 45% and 62% respectively (**Figure 5f**). This is consistent with a previous report demonstrating reduced *elt-3* expression after *pal-1* RNAi as determined by a fluorescent transcriptional reporter (Baugh et al. 2005). The *elt-2* and *elt-3* genes both encode a member of the GATA transcription factor; *elt-2* functions in the intestine to regulate intestinal differentiation while *elt-3* primarily regulates epidermal gene expression (Fukushige et al. 1998; Gilleard & McGhee 2001). Both *elt-2* and *elt-3* have been shown to serve as a co-factor to the SKN-1 transcription factor, with *elt-2* required for *skn-1* dependent pathogen response and *elt-3* required for *skn-1* dependent oxidative stress response (Block et al. 2015; Hu et al. 2017). Consistently, depletion of *elt-2* compromises *C. elegans* immunity and mutation to *elt-3* decreases *C. elegans* resistance to oxidative stress (Hu et al. 2017; Budovskaya et al. 2008; Block & Shapira 2015). We found that knockdown of *pal-1* also significantly reduced *elt-2* and *elt-3* mRNA expression under cadmium (−58%, −60%), and acrylamide (−53%, −65%) conditions in comparison to EV RNAi (**Figure 5g**). The 53% reduction in *elt-2* mRNA under acrylamide condition was not statistically significant (P=0.13), this is perhaps due to the stringency of the two-way ANOVA test used. Together, these results show that the expression of two stress transcriptional co-regulators *elt-2* and *elt-3* are reduced in response to *pal-1* knockdown, implicating a potential mechanism through which depletion of *pal-1* compromises *C. elegans* stress resistance.

Overall, these results show that PAL-1 interacts with CCF-1 protein, and that knockdown of *pal-1* reduces *C. elegans* survival to cadmium and acrylamide stress by potentially attenuating the mRNA expression of the *elt-2* and *elt-3* transcription factors that are required to mount a transcription response to environmental stress (**Figure 5h**).

## DISCUSSION

Transient activation of gene transcription provides a strategy for cells to adapt and respond to stressful conditions based on environmental demand and is critical for organismal survival. Under stress, transcription factors are recruited to gene promoters to initiate transcription based on the distress signal and the response required, and multiple transcription factors often function in parallel for a synergetic response. Here, we show that the *C. elegans* CCR4-NOT subunit CCF-1 physically interacts with the PAL-1 transcription factor and both are required for the transcriptional activation of multiple classes of stress protective genes in response to cadmium and acrylamide. Knockdown of *pal-1* reduces the gene expression of *elt-2* and *elt-3* encoding transcriptional co-regulators that are involved in multiple stress responses (Block & Shapira 2015). Given that *ccf-1* and *pal-1* are required for activating a broad range of genes that are controlled by multiple transcription factors, we propose that these two genes function to positively promote *elt-2* and *elt-3* expression that serves to co-regulate gene transcription during periods of environmental stress (**Figure 5h**).

### CCR4-NOT in stress response and transcription

A role for the CCR4-NOT complex in stress response was initially demonstrated in yeast where CCR4-NOT mutants showed increased sensitivity to replication stress and defects in accumulating *RNR* (ribo<u>n</u>ucleotide reductase) genes in response to DNA damage (Mulder et al. 2005). Mechanistically, it was later shown that CCR4-NOT is recruited to the promoter and open reading frames of *RNR* genes to facilitate transcription initiation and elongation in response to DNA damage (Kruk et al. 2011). Genome-wide crosslinking analyses have also shown that CCR4-NOT is predominantly recruited to SAGA-regulated genes in yeast that typically encode highly inducible genes, suggesting a potential role for CCR4-NOT in stress regulation in yeast (Venters et al. 2011). In human cells, CCR4 can serve as a coactivator of several nuclear hormone receptors where CCR4 overexpression enhanced the activation of these receptors while CCR4 knockdown reduced receptor activation and decreased the abundance of hormone responsive target genes (Garapaty et al. 2008). In mice, the CCF-1 homolog CNOT7 directly interacts with retinoid X receptor β to function as a transcriptional co-activator and the knockout of *Cnot7* impairs retinoic acid induced transcription (Nakamura et al. 2004). We show in this study that multiple subunits of the CCR4-NOT complex positively promotes transcription of stress inducible genes in *C. elegans*. In this instance, the CCF-1 subunit of the CCR4-NOT complex interacts physically and functionally with the PAL-1 transcription factor in promoting stress induced gene expression. Knockdown of *ccf-1* compromises the oxidative stress resistance of *C. elegans* as evident by reduced survival to cadmium and acrylamide stress along with increased sensitivity to acrylamide induced dopaminergic neurodegeneration. Given that both *ccf-1* and *pal-1* knockdown affected a broad network of stress responsive gene classes, we propose this interaction likely indirectly promotes stress resistance by activating the expression of *elt-2* and *elt-3* that encode master transcriptional co-regulators involved in multiple stress responses (Block & Shapira 2015). It remains to be determined whether PAL-1 interacts with the CCR4-NOT complex as a whole or just CCF-1 alone. However, our results showed that knockdown of other CCR4-NOT subunits including *ntl-2, ntl-3*, and *let-711* can also attenuate activation of cadmium induced genes, suggesting the involvement of the CCR4-NOT complex as a whole in environmental stress response.

### The elt transcriptional network

The GATA transcription factors *elt-2* and *elt-3* have both been shown to function with transcription factors in the *C. elegans* intestine and epidermis respectively to regulate the expression of a wide network of stress responsive genes. Following infection to *Pseudomonas aeruginosa*, ELT-2 cooperates with the transcription factors SKN-1 and ATF-7 (<u>A</u>ctivating <u>T</u>ranscription <u>F</u>actor) to promote the expression of different classes of immune genes (Block & Shapira 2015). ELT-2 functions as the predominant regulator of intestinal gene expression, which is the major site of pathogen and xenobiotic stress response in *C. elegans.* Consistently, candidate direct targets of ELT-2 predicted through SAGE analysis include those functioning in metal detoxification, cytochrome P450 metabolism, and glutathione activity (McGhee et al. 2009). Analogous to ELT-2, ELT-3 appears to also function as a stress transcriptional co-regulator with the distinction that ELT-3 primarily acts in the epidermis (Block & Shapira 2015). In response to oxidative stress, ELT-3 directly binds to the SKN-1 transcription factor to promote the expression of *gst* related antioxidant genes (Hu et al. 2017). ChIP analysis indicates that one-third of ELT-3’s targets are also bound directly by the SKN-1 transcription factor, reinforcing the role of ELT-3 as a co-regulator of SKN-1 (Block & Shapira 2015). In response to pathogen infection and osmotic stress, ELT-3 controls the expression of neuropeptide-like protein genes potentially through a mechanism involving the STA-2 (<u>S</u>ignal <u>T</u>ransducer and <u>A</u>ctivator) transcription factor (Pujol et al. 2008; Dodd et al. 2018). Lastly, ELT-3 is also required for the expression of DAF-16 regulated genes in the hypodermis, suggesting that it also functions within the insulin/IGF-1 pathway (Zhang et al. 2013).

We have previously shown that oxidative stress promotes the nuclear localization of SKN-1::GFP in both the intestine and the epidermis, and this is supported by the up-regulation of the *gst-4p*::GFP reporter in both tissues (Wu et al. 2016). Knockdown of *skn-1* in either the intestine or epidermis alone reduces *C. elegans* resistance to oxidative stress, supporting the idea that SKN-1 activity in both tissues is critical for oxidative stress resistance (Wu et al. 2016). Given that our data show that knockdown of *pal-1* reduces both *elt-2* and *elt-3* transcript levels under cadmium and acrylamide stress and that both GATA transcription factors are directly implicated in regulating a network of stress protective genes, it is conceivable that *pal-1* can broadly influence stress response pathways indirectly via its transcriptional control over *elt-2* and *elt-3.* Consistent with this, our data show that knockdown of *pal-1* decreases *C. elegans* resistance to cadmium and acrylamide stress. However, we cannot rule out the possibility that PAL-1 may also directly bind to the promoters and initiate the transcription of the target genes in response to stress.

### Role of ccf-1 in aging

Genes that regulate stress response in *C. elegans* are often also factors that influence aging. Our results show that knockdown of *ccf-1* shortens normal lifespan as well as the longevity of *skn-1, hsf-1*, and *daf-16* gain of function or overexpression mutants. In the *skn-1* gain of function mutant, *ccf-1* was completely required for its longevity effect and this is consistent with our transcriptome analysis revealing that *ccf-1* dependent stress genes highly cluster to glutathione metabolism. Interestingly in the *daf-2* mutant where *daf-16* is activated, the relative lifespan extension compared to N2 was similar when worms were fed with EV or *ccf-1* RNAi. This could be inferred as that the mechanism through which *ccf-1* knockdown shortens lifespan is independent of the downstream pathways through which the *daf-2* mutant extend longevity. While this interpretation would conflict with evidence that *skn-1* functions downstream of *daf-2* mutant longevity, a possible explanation is that knockdown of *ccf-1* may not restrict other regulatory functions of *skn-1* such as lipid metabolism or collagen gene expression that have been directly implicated in longevity regulation (Steinbaugh et al. 2015; Ewald et al. 2015). In support of a diminished role for *skn-1* dependent xenobiotic genes in *daf-2* mutant longevity, we previously showed that loss of function mutation to the *xrep-4* (<u>x</u>enobiotics <u>re</u>sponse <u>p</u>athways) gene that specifically attenuates *skn-1* dependent *gst* gene expression does not reduce wildtype or *daf-2(e1037)* lifespan (Wu et al. 2017). It is also possible that given the pleiotropic function of the CCR4-NOT complex in mRNA regulation, its role in lifespan regulation may be linked to its deadenylase activity not explored in this study. Nonetheless, these data clearly demonstrate a role for the CCR4-NOT complex in lifespan regulation and would be a subject of interest in future studies.

Overall, our study provides evidence that the CCR4-NOT complex is a key regulator for the transcriptional response to various environmental stressors in *C. elegans*. Given the broad range of protective genes regulated by *ccf-1*, we propose that this mechanism signals through the PAL-1 transcription factor to indirectly influence stress resistance by regulating the gene expression of the *elt-2* and *elt-3* tissue specific transcriptional co-regulators.

## MATERIALS AND METHODS

### C. elegans strains

The following strains were used: N2 Bristol wildtype, QV151 *qvIs4[numr-1p::GFP]*, BZ555 *egIs1[dat-1p::GFP]*, VP198 *kbIs5[gpdh-1p::GFP; rol-6(su1006)]*, QV212 *skn-1(k1023)*, AGD710 *uthIs235 [sur-5p::hsf-1::unc-54 3’UTR; myo-2p::tdTomato::unc-54 3’ UTR]*, CB1370 *daf-2(e1370) III*, MWU110 *cwwIs4 [ccf-1p::*CCF-1*::GFP::unc-54 3’UTR; myo-2p::tdTomato::unc-54 3’ UTR]*, NL2099 *rrf-3(pk1426) II.* All *C. elegans* strains were cultured at 20 °C using standard methods (Brenner 1974), with the exception of *daf-2(e1037)* and *rrf-3(pk1426)* which were grown at 16 °C during development followed by shift to 20 °C on the first day of adulthood.

### Genome-wide RNAi screen and RNAi experiments

RNAi screen was performed as previously described in detail (Wu et al. 2019). Briefly, synchronized L1 QV151 larvae were obtained using hypochlorite treatment and grown in liquid nematode growth media (NGM) and fed with dsRNA producing HT115(DE3) bacteria for 2 days, followed by exposure to 100 μM cadmium chloride for 24 h and screened for suppression of *numr-1p*::GFP fluorescence. Approximately 19,000 dsRNA clones from the MRC genomic RNAi feeding library (Geneservice, Cambridge, UK) and the ORFeome RNAi feeding library (Open Biosystems, Huntsville, AL) were screened. Clones that suppressed *numr-1p*::GFP fluorescence were rescreened three additional times for validation. For all other RNAi experiments in this study, NGM agar plates containing 50 μg mL^−1^ carbenicillin and 100 μg mL^−1^ of isopropyl β-D-thiogalactopyranoside (IPTG) were used. *E. coli* expressing the pPD129.36 (LH4400) plasmid was used as an RNAi control and referred to as empty vector (EV) in this study as this plasmid encodes 202 bases of dsRNA not homologous to any *C. elegans* genes. For the *pal-1* RNAi experiments, the enhanced RNAi sensitive strain *rrf-3 (pk1426)* was used.

### Microscopy and fluorescent analysis

To analyze *numr-1p*::GFP and *gpdh-1p*::GFP transcriptional reporters or the CCF-1::GFP translation reporter, synchronized worms were grown on EV or *ccf-1* RNAi seeded NGM agar plates until day 1 of adulthood followed by transfer to NGM agar plates seeded with corresponding RNAi containing 100 µM of cadmium for *numr-1p*::GFP, 300 mM NaCl for *gpdh-1p*::GFP, and 100 µM of cadmium or 5 mM of acrylamide for CCF-1::GFP analysis. After 24 hours, worms were mounted on a glass slide containing a 2% agarose pad and immobilized with 2% sodium azide before imaging with a Zeiss Axioskop 50 microscope mounted with a Retiga R3 camera. Eight worms were mounted per slide and relative fluorescence was calculated using the Measure function in ImageJ followed by background subtraction. The background signal for each image was calculated by defining an area on the same image where the fluorescence signal was absent, with the dimension of the background area subtracted constant for all images.

Methods used to assess acrylamide induced dopaminergic degeneration via *dat-1p*::GFP were described previously in detail (Murray et al. 2020). Briefly, synchronized L1 *C. elegans* were grown on NGM agar plates seeded with EV or *ccf-1* RNAi until day 1 of adulthood followed by transfer to NGM agar plates containing 5 mM of acrylamide seeded with the corresponding RNAi bacteria and grown until 6 days old. Adult worms were separated from their progeny via picking with a sterilized metal pick onto fresh acrylamide agar plates during the reproductive window. On day 6, worms were prepared for imaging using the procedures described above.

Scoring criteria for dopaminergic integrity were as follows, wildtype indicates smooth and continuous cephalic neuron (CEP) sensilla dendrite located at the anterior portion of the worm, blebbing indicates at least 1 abnormal punctae along the CEP dendrites, and breaks indicate discontinuation within the CEP dendrite. Three trials were performed with N = 15 worms scored per condition each trial. All grayscale images were converted to color using ImageJ, with composite images displaying both GFP and RFP colors created using the Merge Channel function when applicable.

### RNA extraction and qPCR

The Purelink RNA mini kit (ThermoFisher, 12183020) was used to isolate total RNA with a QSonica Q55 sonicator used for tissue lysis. Synchronized L1 N2 or *rrf-3(pk1426)* worms were grown on NGM agar plates seeded with EV or corresponding RNAi until day 1 of adulthood followed by transfer to either control agar plates, agar plates containing 100 μM of cadmium chloride, or agar plates containing 5 mM of acrylamide, each seeded with the corresponding RNAi bacteria for an additional 24 hours followed by RNA extraction. For the heat shock experiment, synchronized L1 N2 worms were fed with EV or *ccf-1* RNAi until day 1 of adulthood followed by heat shock at 37 °C for 1 hour followed by RNA extraction. For each condition, N = 4 biological RNA replicates were prepared with each replicate containing ~ 500 worms. For qPCR, RNA was first treated with DNAseI (ThermoFisher, EN0521) followed by cDNA synthesis with the Invitrogen Multiscribe^TM^ reverse transcriptase (ThermoFisher, 4311235) using an Applied Biosystems ProFlex thermocycler. Real-time PCR (qPCR) was performed with the PowerUp^TM^ SYBR^TM^ Green Master Mix (#A25741) in a QuantStudio3 system. Relative gene expression was normalized to the housekeeping gene *rpl-2* (ribosomal <u>p</u>rotein large subunit) and primers used for this study are shown in Table S3.

### RNA-sequencing and data analysis

Wildype N2 worms synchronized at the L1 stage were grown on NGM agar plates seeded with EV or *ccf-1* RNAi until day 1 of adulthood followed by transfer to either control agar plates, agar plates containing 100 μM of cadmium chloride, or agar plates containing 5 mM of acrylamide, each seeded with the corresponding EV or *ccf-1* RNAi for 24 hours followed by RNA extraction. Total RNA from three biological replicates for each experimental group was extracted using the methods described above, with the exception that ~2,000 to 3,000 worms were used for each replicate. RNA samples were then sent to Novogen (Sacrament, CA) on dry ice for cDNA library construction with oligo(dT) enrichment and sequencing. Sequence annotation and data analysis were performed by Novogene.

### Lifespan and survival assays

All assays were performed at 20°C using methods previously described (Wu et al. 2019), except for *daf-2(e1037)* and *rrf-3(pk1426)* which were grown at 16°C during development. For lifespan assays, synchronized L1 worms obtained through the hypochlorite treatment were grown on the appropriate RNAi seeded NGM agar plates until adulthood. One day old adult worms were moved to new plates and continuously transferred via daily picking during the reproductive window to accomplish progeny separation. For survival assays, synchronized L1 worms were first grown on control NGM plates seeded with appropriate RNAi bacteria until the first day of adulthood, followed by transfer to NGM plates containing either 100 μM of cadmium chloride or 5 mM of acrylamide seeded with the corresponding RNAi bacteria. For both lifespan and survival assays, the first day of adulthood is considered as 1 day old, and worms were scored every 1-2 days for death by gentle prodding with a flame sterilized metal pick. Worms that do not respond to the gentle prodding were considered dead, and worms with protruding vulva or gonad were recorded as censors. Three independent trials were formed for all assays with the number of animals scored in each condition and experiment listed in Table S2.

### Y2H library screen

The full-length *ccf-1* cDNA clone was generated through PCR using the Q5® High-Fidelity DNA polymerase (New England BioLabs, M0491L) and cloned into the pDEST32 vector containing the GAL4 DNA binding domain (DBD) through Gateway cloning (ThermoFisher, 11789020 and 12538120) and used as the Y2H bait construct. A Y2H prey cDNA library of *C. elegans* genes was cloned into the pDEST22 vector containing the GAL4 activating domain (AD) using the CloneMiner^TM^ cDNA library construction kit (ThermoFisher, A11180). A forward Y2H library screen was performed using the ProQuest Two-Hybrid System (ThermoFisher, PQ1000101) to identify protein interactors of the *C. elegans* CCF-1 protein.

Yeast colonies containing bait and prey constructs that grew on Trp^−^/Leu^−^/His^−^ + 25 mM 3-Amino-1,2,4-triazole (3AT) selection plates were extracted using the Zymoprep Yeast Plasmid Miniprep II (Zymo Research, D2004) followed by Sanger sequencing for gene identification. Three independent colonies from each bait-interacting prey construct were further tested for its interaction with CCF-1 by evaluating for 1) growth on Trp^−^/Leu^−^/Ura^−^ selection plate, 2) negative growth on Trp^−^/Leu^−^ + 0.2% 5-fluoroorotic acid selection plate, 3) growth on Trp^−^/Leu^−^/His^−^ + 25 mM AT selection plate, and 4) positive phenotype from the X-gal assay. Yeast cells were normalized to OD_600_ = 0.5 for the 10^0^ concentration followed by serial dilution on the growth assay. Prey construct that passed 3 out of 4 selection tests was considered as a positive CCF-1 interacting protein. Yeast cells were imaged on a Bio Rad Gel Doc EQ system.

### Statistical analyses

Graphical data and statistical analysis were performed using the Graphpad Prism software (V7.04). Student’s t-test was used for comparison of two groups with the Holm-Sidak multiple test correction applied when the test is repeated for multiple genes, one-way ANOVA with Dunnett’s test was used for comparison of more than two groups, two-way ANOVA with Holm-Sidak test was used for comparison of two factors, categorical data were analyzed using the Chi-square test, and linear-regression was analyzed using the F-test. Lifespan and survival data were analyzed with the log-rank test using the OASIS2 software (https://sbi.postech.ac.kr/oasis2/history/). For RNA-sequencing data, false discovery rate (FDR) correction was applied to determine the statistical significance. For all statistical tests, the following designations were used to indicate P-value, *P<0.05, **P<0.01, ***P<0.001.

## ACKNOWLEDGEMENTS

Some *C. elegans* strains were provided by the *Caenorhabditis* Genetic Centre (University of Minnesota, Minneapolis, MN) which is supported by the NIH Office of Research Infrastructure Programs (P40 OD010440). We thank Dr. Carlos Carvalho (University of Saskatchewan) for discussion on the Y2H screen. CWW is supported by an NSERC Discovery Grant (04486), KPC was supported by NSF grants IOS-1120130 and IOS-1452948, HT and BMW were supported by a WCVM graduate teaching fellowship and devolved scholarship respectively, and KR was supported by an NSERC USRA scholarship.

## AUTHOR CONTRIBUTIONS

HT, BMW, KR, and SMM performed experiments and analyzed the results. KPC helped conceive and design the RNAi screen; YW and KPC generated the Y2H cDNA library. CWW designed the study, performed experiments, analyzed results, and wrote the manuscript.

## DATA AVAILABILITY

All datasets supporting this manuscript are found within the article and its supplementary files. RNA-sequencing data generated from this study (raw and annotated) are available on the NCBI GEO data repository GSE194057. Strains are available upon request.

## SUPPORTING INFORMATION

**Figure S1.**
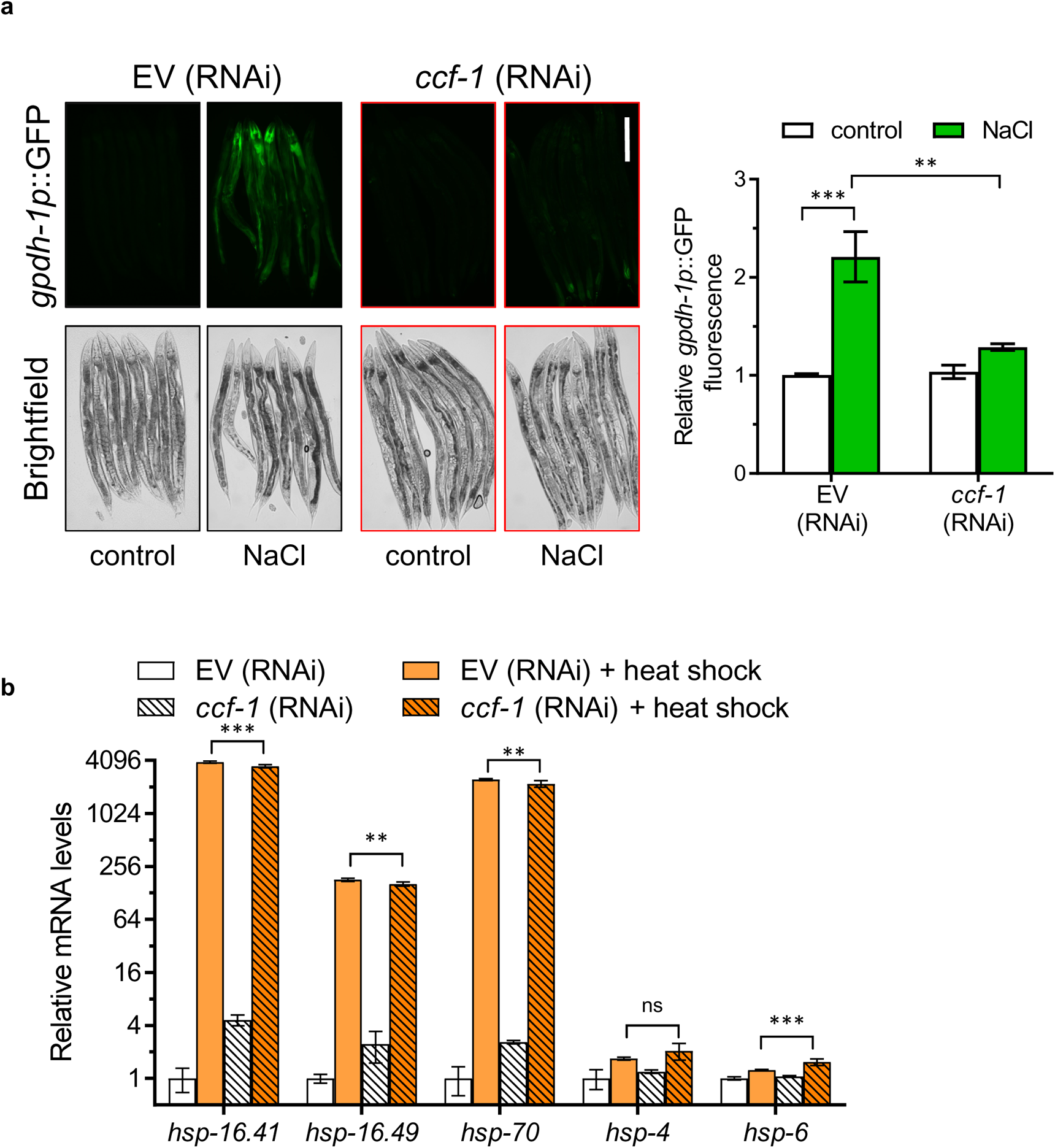
Knockdown of *ccf-1* attenuates the osmotic and heat stress response. **a.** Relative fluorescence level of *gpdh-1p*::GFP in worms fed with EV or *ccf-1* (RNAi) under basal conditions or exposed to NaCl. Shown on the right are representative fluorescent and brightfield micrographs. Scale bar = 100 µm. **b.** Relative expression of *hsp* genes in worms fed with EV or *ccf-1* (RNAi) under control conditions or heat shock. *P<0.05, **P<0.01, and ***P<0.001 as determined by two-way ANOVA with Holm-Sidak *post hoc* tests.

**Figure S2.**
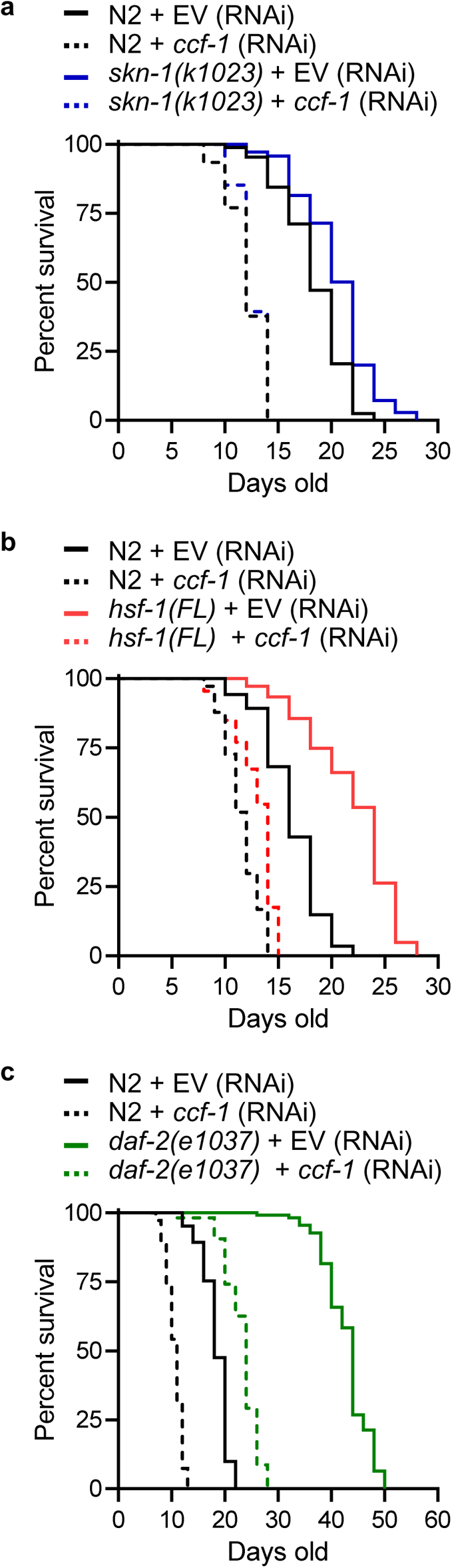
*ccf-1* is required for the longevity of long-lived mutants. Lifespan curves of N2 and **a.** *skn-1(k1023),* **b.** *hsf-1(FL)*, and **c.** *daf-2(e1037)* worms fed with EV or *ccf-1* (RNAi). Results for trial 1 are shown for all assays with full results and statistics for all trials are presented in Table S2.

**Figure S3.**
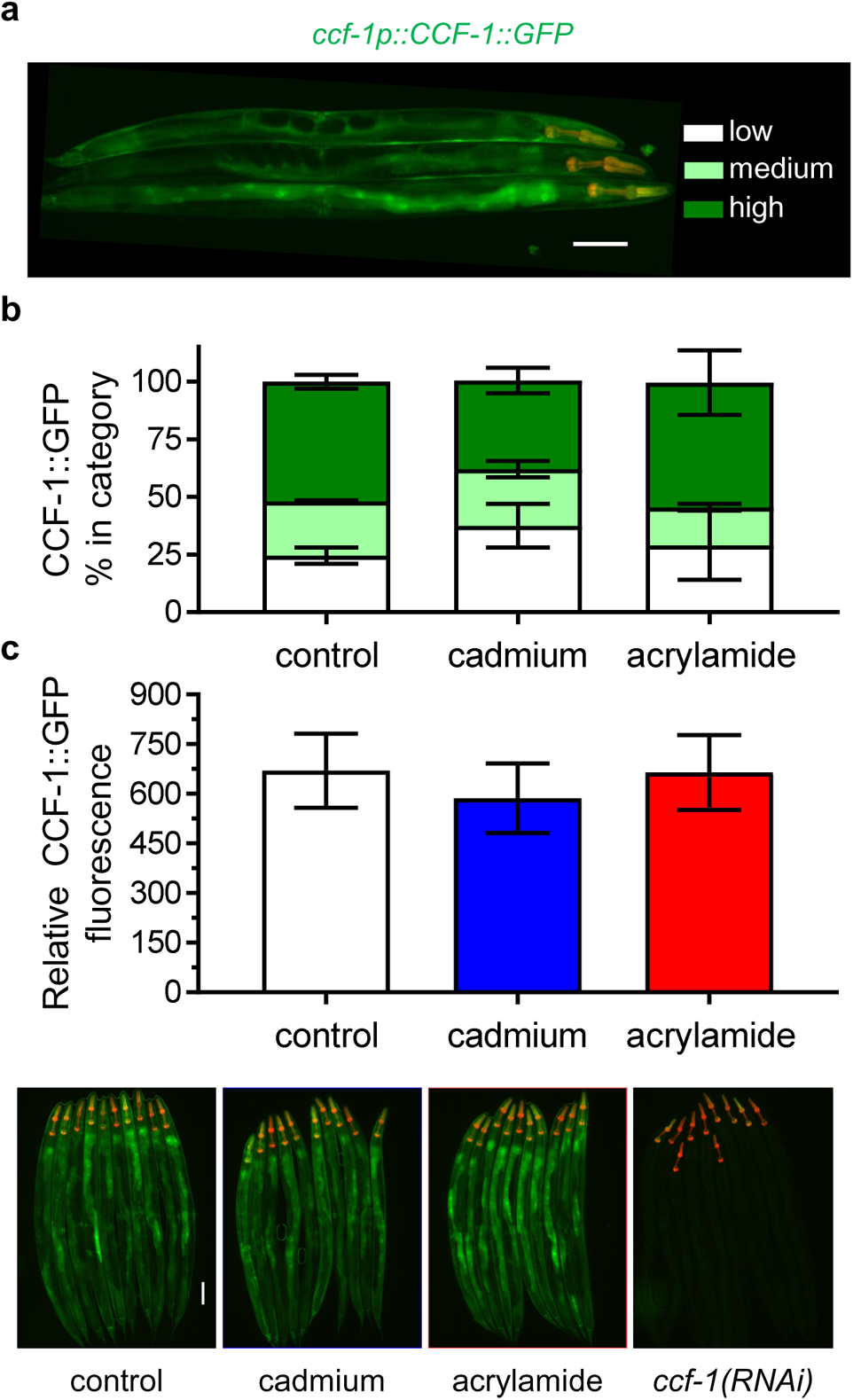
CCF-1::GFP expression is not affected by stress. **a.** Representative fluorescent micrographs showing low, medium, and high levels of nuclear CCF-1::GFP localization. Scale bar = 100 µm. Relative **b.** nuclear distribution and **c.** fluorescence of CCF-1::GFP in worms under control, cadmium, and acrylamide exposure conditions. Representative fluorescence micrographs are shown below for each condition along with control worms fed with *ccf-1* (RNAi). Results are combined from two trials of N = 90-100 worms per condition scored each trial.

**Figure S4.**
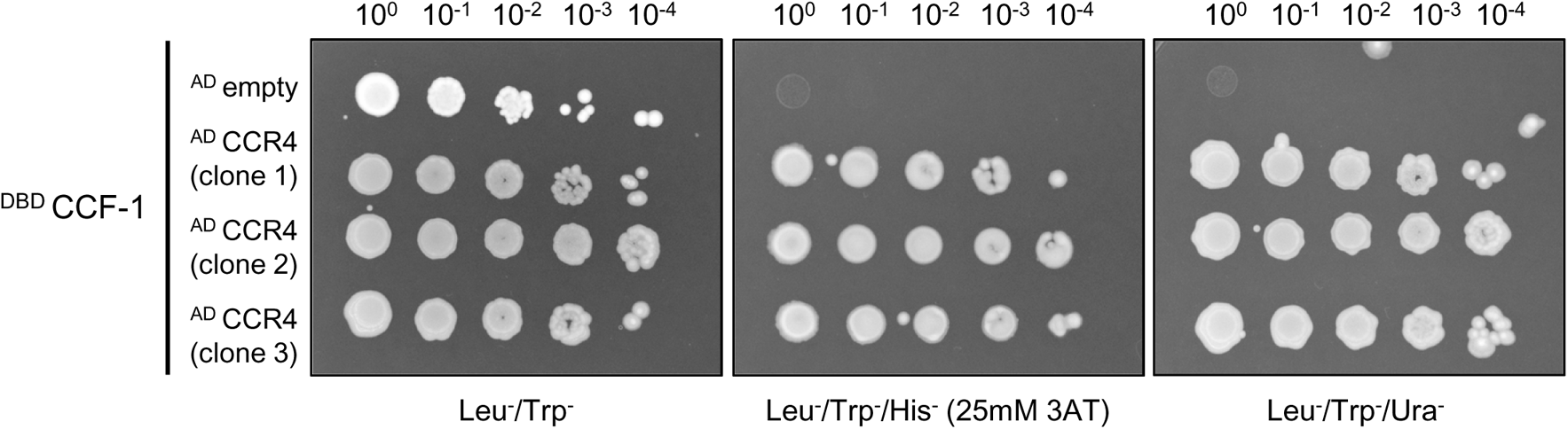
Yeast two-hybrid interactions between CCF-1 (bait) and three independent clones of CCR4 (prey) across a dilution gradient on the control Leu^−^/Trp^−^ plate, and Leu^−^/Trp^−^/His^−^/25 mM 3AT or Leu^−^/Trp^−^/Ura^−^ selection plates.

**Table S1.** Genome-wide RNAi list

**Table S2.** Lifespan data summary

**Table S3.** qPCR primers used in this study

